# When and how leachate toxicity of tire wear particles peaks: quantifying its dynamics using dose-response analysis

**DOI:** 10.1101/2024.03.30.587449

**Authors:** Yanchao Chai, Xin Wang, Haiqing Wang, Yu Zhang, Jiaxin Yang

## Abstract

The aquatic toxicity of tire wear particles (TWPs) depends on the leaching of various components. The leaching process is time-dependent and influenced by various environmental factors, leading to fluctuation in leachate toxicity. In this study, we quantified the temporal dynamics of leachate toxicity using the dose-response model. Meanwhile, to simulate weathering conditions, TWPs were exposed to ultraviolet (UV) irradiation and elevated temperature. The results indicated that the leachate toxicity over time fitted a biphasic pattern, with an initial increase followed by a decrease, peaking at the 11^th^ day. The toxicity fluctuation cannot be characterized by any single component, but rather leachate should be considered as an integral whole aligning with a typical hormesis dose-response curve. High temperature of road surface increased leachate toxicity by 66%, due to elevated concentrations of 67 identified organic compounds. UV irradiation, however, mitigated this effect, acting as antagonism. These findings underline the seasonal and spatial heterogeneity in leachate toxicity. Additionally, high temperature induced the volatilization of organic compounds within TWPs, highlighting an independent exposure pathway. The methodology developed in this study will provide foundation and standard for future research on the aquatic ecotoxicity of TWPs.

**Environmental Implication:** The toxicity of leachate from ubiquitous tire wear particles (TWPs) to aquatic organisms has been confirmed in field and experimental studies. However, existing researches indirectly used TWPs quantity as sole proxy for leachate toxicity, neglecting its fluctuations over leaching duration and in situ environmental factors, potentially confounding toxic effects. Herein, this study quantifies these dynamics of leachate toxicity based on measurable biological response, aiming to establish a standard scale for further quantitative assessment of TWPs ecological risk.

## 1. Introduction

The leachate of tire wear particles (TWPs) has been demonstrated to pose significant threats to numerous aquatic organisms in both controlled toxicological experiments and field studies. (Capolupo et al., 2021; Khan et al., 2019; Marwood et al., 2011; Panko et al., 2013; Tian et al., 2021; Turner and Rice, 2010; Wik and Dave, 2006). The ecotoxicological effects of tire wear particles (TWPs) are largely a consequence of the leaching of numerous additives and their transformation products present in the tire formulation. (Boisseaux et al., 2024; Li et al., 2024). A quantitative assessment of leachate toxicity necessitates a well-defined dose metric. In prior investigations, the mass or concentration of tire wear particles (TWPs) has served as a proxy for leachate dose. However, upon the introduction of TWPs into aquatic environments, concurrent processes such as the release and biodegradation of toxic constituents occur, rendering leachate toxicity temporally dynamic. Consequently, leachate derived from identical masses of TWPs may exhibit varying toxic potencies at different time points. Inconsistent extraction times, spanning from a few hours to multiple weeks, used in the preparation of TWP leachate across studies pose a significant methodological issue. This variability, coupled with the temporal fluctuations in leachate toxicity, can lead to discrepancies and confusion. Hence, quantitative characterization of leachate toxicity is warranted, moving beyond the simplistic representation by TWP mass.

In theory, leachate toxicity can be directly characterized by quantifying the concentrations of its leachable components. Specific components identified as highly toxic in previous studies, exhibit concentration changes over leaching time, necessitating a dynamic assessment of their potential ecological impact (Foscari et al., 2024; Li et al., 2023). However, this approach is limited as it fails to account for the complex interactions among the hundreds of components in TWP leachate (Li et al., 2024). The dose-response curve, a fundamental tool in pharmacology and toxicology, quantifies the relationship between effector dose and biological response, aiding in the understanding of the potency and efficacy of various substances (Calabrese and Baldwin, 2003; Waddell, 2010). Thus, leachate toxicity can be characterized and quantified through biological response. Subsequently, its time-dependent dynamics will be fitted using a kinetic model.

Besides the leaching time, it has been shown that certain weathering parameters, such as temperature, ultraviolet (UV) irradiation, abrasion, and oxidation, can alter the biological toxicity of TWPs and microplastics (Kolomijeca et al., 2020; Lv et al., 2024; Zhang et al., 2024). Prior to entering aquatic environments, TWPs generated on road surfaces undergo a migration process during which they are exposed to various environmental factors. During summer months, for instance, road surface temperatures can reach 70℃ coupled with high levels of UV irradiation. The weathering conditions experienced by *in situ* TWPs are frequently overlooked during their artificial production for experimental use, potentially resulting in inaccurate assessments of leachate toxicity. Given daily urban road washing, exposure within one day to high temperature and UV irradiation is environmentally realistic. The spatiotemporal variability of these weathering conditions implies regional and seasonal differences in TWP leachate toxicity, which are crucial for refining TWPs leachate risk assessments. This enables the development of targeted policies to minimize ecotoxicity across varying regions, seasons, and weather conditions. Moreover, tire additives with lower boiling points can volatilize into the air under high temperatures (Johannessen et al., 2022; Kim and Lee, 2018), representing a potential novel exposure pathway independent of TWPs that warrants further investigation.

To quantify the dynamic leachate toxicity of TWPs over time, we used biological response as the dependent variable in a dose-response analysis. The freshwater model zooplankton *Brachionus calyciflorus* was employed as the indicator organism, with reproduction inhibition as the endpoint due to its high sensitivity to TWPs leachate. (Chai et al., 2025). Meanwhile, the influence of weather conditions, such as UV irradiation and high temperature, on the leaching toxicity of TWPs was characterized as *in situ* modification. Furthermore, the potential volatilization of additives in TWPs under high temperatures was investigated as a novel exposure pathway. This research is likely to contribute to the standardization of ecological risk assessments for TWPs leachate.

## 2. Materials and Methods

### 2.1 TWPs and rotifer

TWPs were purchased from a factory that recycles and smashes scrap tires into particles as artificial turf material. These TWPs were sieved by turn through 100 μm and 38 μm nylon net, and the fraction between these two sizes was collected and stored at 4℃.

Rotifer *B. calyciflorus* clone strain was established from resting egg that was provided by Professor T.W. Snell (Georgia Institute of Technology, USA). The hard synthetic freshwater was used as culture medium (96 mg NaHCO_3_, 60 mg CaSO_4_, 60 mg MgSO_4_ and 4 mg KCl in 1 L distilled water) (USEPA, 2002). The culture was maintained in illumination incubator with 16L:8D light schedule with light intensity of 4000 Lux at 25℃. Rotifers were fed with 5×10^6^ cell/ml *Chlorella pyrenoidesa*, and the medium was refreshed daily.

### 2.2 Preparation of TWPs leachate at different extraction times

The soaking time points were 3, 7, 15, 30, and 60 days. For each treatment, TWPs were soaked in 2.5 L of hard synthetic freshwater (2500 mg/L) in a glass bottle and mixed. The mixture was continuously aerated in the dark at 25℃. At each time point, the leachate stock solution was obtained by filtering the mixture through a 0.45 μm acetate fiber filter and stored at -20℃.

### 2.3 Exposure of high temperature and UV radiation to TWPs

Four treatments were established: control (CK), high temperature (Heat), UVB radiation (UV), and sequential high temperature and UVB radiation (Heat+UV). TWPs in glass Petri dishes were exposed to 70℃ for 12 hours, mimicking the temperature of asphalt pavement in summer. This was followed by 11 hours of UVB radiation at an intensity of 2 mW/cm^2^ (total dose: 81300 mJ/cm^2^, simulating the daily irradiation at 32°N latitude during summer). The samples were stirred every two hours during UVB exposure to ensure uniformity. Following these pretreatments, TWPs were soaked to prepare stock solutions as detailed in Section 2.2.

### 2.3 Collection of soluble components from volatilization

A 500 mL glass gas-washing bottle containing 2.5 g of TWPs was sealed and placed in a 70℃-thermostat water bath for 12 hours. The inlet was connected to an air pump with a flow rate of approximately 6 L/min, simulating human breathing rate. The exhaust, filtered through a 0.45μm filter, was introduced to the bottom of another gas-washing bottle containing 500 mL of deionized water. The solution containing soluble volatile substances was frozen at -20℃ for component analysis.

### 2.4 Toxicity test of different TWPs leachates on rotifer

The TWPs leachate stock solution from every treatment was diluted by rotifer culture medium with concentration gradient (0, 250, 500, 750, 1000 mg/L). Rotifer with amictic egg were selected from clone strain and synchronized (Kaneko et al., 2016). Then neonate born within 4 hours were put into 24-well plate. There was one individual in one well with 1 ml the specific concentration TWPs leachate, in which algae were added at the density of 5×10^6^ cell/ml. The plates were placed in illumination incubator with 16L:8D light schedule with light intensity of 4000 Lux at 25℃. The neonates were checked and picked out every 12 hours till all individuals died. Each treatment included 48 individuals. The fecundity (offspring number) was regarded as the indicator of biological response.

### 2.5 Detection of components in leachate

The relative quantitative analysis of organic compounds in leachate was determined by HPLC-MS (Thermo, Ultimate 3000LC, Q Exactive HF). The detailed conditions were showed in Supplementary Information (Section 1, Table S1). According to the peak area of metabolites in each sample, the relative concentrations (μg/mL) in the samples were calculated.

The concentration of heavy metal elements was analyzed by ICP-OES optima 8000 (Perkin Elmer, Co., Ltd., USA) including S、Zn、Sb、Mn、Sr、Co、Cd、Ni、Cr、 Cu. The water samples were digested with nitric acid in a microwave digestion apparatus (TOPEX) before analysis.

### 2.6 Statistics

The average individual fecundity of rotifers in each treatment was calculated by dividing the total offspring number by the number of individuals, excluding mictic individuals. The biological response was expressed as the relative value (%) of fecundity compared to the control group (0 mg/L). (Formula 1). Positive value means promotion effect, otherwise inhibition.

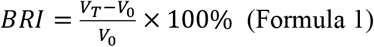

Where BRI was biological response index, V_T_ and V_0_ was in treatment group and average offspring number control group.

The Person correlation analysis was executed by IBM SPSS Statistics version 22, and curve fitting was conducted in GraphPad Prism version 9.5.0.

## 3. Results and Discussion

### 3.1 Quantifying dynamic of leachate toxicity over leaching time

The leachate at different time points exhibited various dose-response relationships (**Figure 1**), showing that the toxicity leached from TWPs is time-dependent. Furthermore, the leachate toxicity conformed to a distinct hormesis dose-response model, characterized by low-dose stimulation and high-dose inhibition, commonly observed in pharmacology (Lushchak, 2014; Sun et al., 2021), which was illustrated in a schematic (**Figure 1**, middle). By comparing the dose-response relationships of the leachate at different extraction times with the typical hormesis curve, we found that these five dose ranges correspond well to different segments of the dose axis (five distinct regions illustrated in **Figure 1**). These results indicate that the toxicity of the leachates at different times can be ranked as follows: 15 days > 7 days > 30 days > 3 days > 60 days. This pattern can be attributed to the content of toxic substances in the leachate being governed by biphasic processes, initially dominated by release from TWPs and subsequently by degradation or sorption. (Foscari et al., 2024).

**Figure 1.**
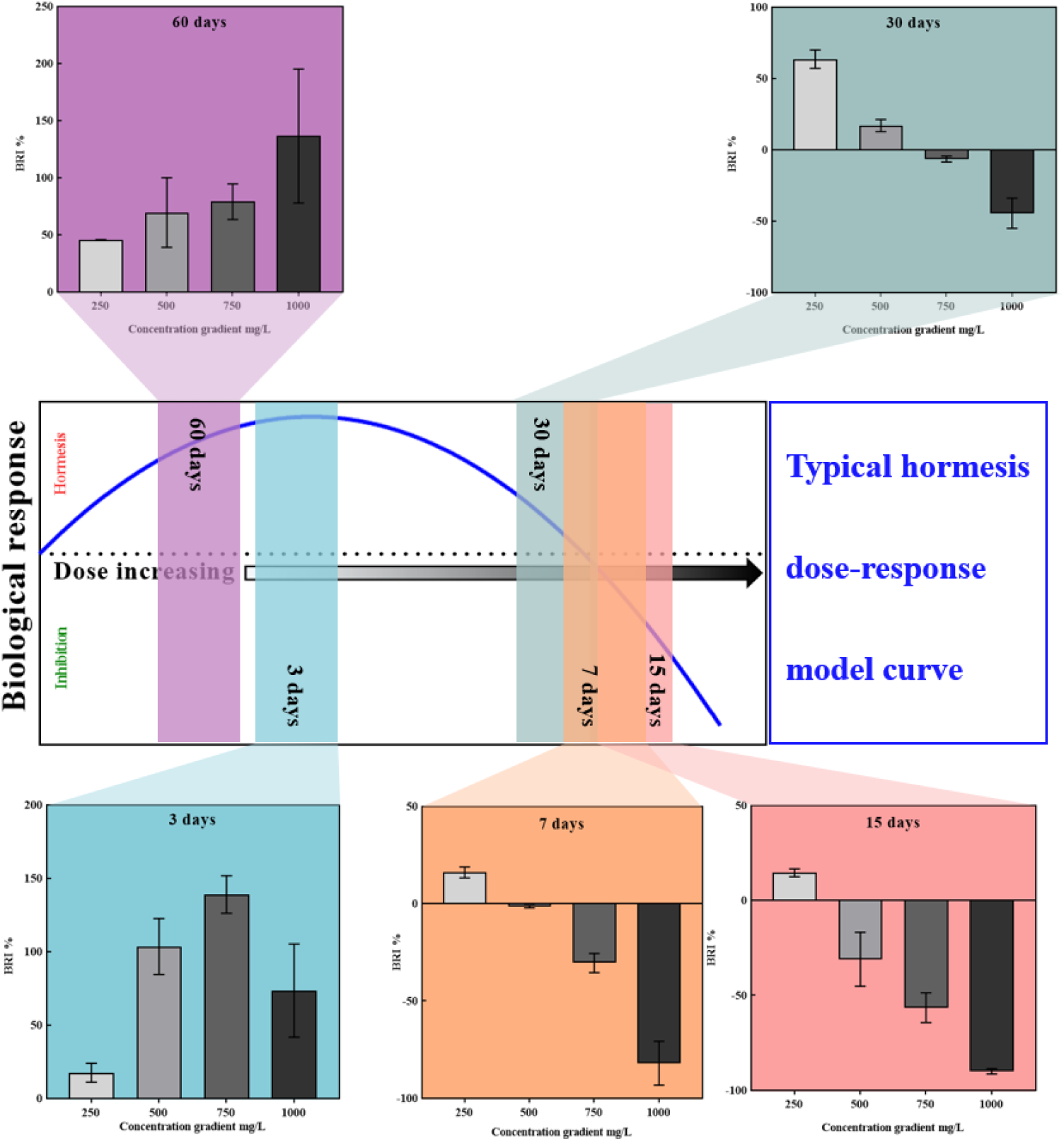
Schematic of typical hermetic dose-response curve and biological responses to TWPs leachate with concentration gradient (250, 500, 500 and 1000 mg/L) from different extraction times of 3, 7, 15, 30 and 60 days. Positive BRI value means hermetic effect, otherwise inhibition; The colorful swaths in Schematic represent relative dose range of leachate toxicity from different extraction times.

To quantify the dynamic of leachate toxicity over time, it is necessary to assess the relative toxicity level at the five different time points. In the hormesis dose-response model, the transition point from stimulation to inhibition is defined as the Zero Equivalent Point (ZEP) (Lushchak, 2014), namely, the right intersection with the x-axis in schematic of **Figure 1**. As for the dose-response relationships of the leachates at the five different time points, their ZEP doses were calculated by regression analysis as follows: 3 days: 1188±87 mg/L; 7 days: 432±24 mg/L; 15 days: 321±16 mg/L; 30 days: 712±24 mg/L; 60 days: 2080±27 mg/L. These ZEP doses were expressed within their respective dose range scales (250-1000 mg/L), referred to as nominal doses, which were represented by the TWPs mass but did not reflect the leachate toxicity. Theoretically, these five nominal ZEP doses correspond to equal toxicity level. A higher nominal ZEP dose indicates that a greater volume of the stock leachate solution is required to achieve the same toxicity level. Therefore, the toxicity of leachate at different extraction times is inversely proportional to its nominal ZEP dose. Then, the dynamics of leachate toxicity level over time based on biological response was quantified, as shown in **Figure 2**. The two-phase curves for were fitted with different functions, specifically quadratic and exponential equations, and their point of intersection was calculated to indicate that leaching toxicity peaked around 11^th^ day. To verify accuracy of this model, we prepared again the leachate after 11-days extraction, and its actual relative toxicity level calculated based on dose-response curve was approaching to the predicted value (**Figure 2**). The first phase reflected the rapid release and accumulation of additives in the TWPs. Subsequently, the second phase was primarily influenced by biodegradation. This trend was corroborated by the changes in concentration of certain components leached from the TWPs (Foscari et al., 2024).

**Figure 2.**
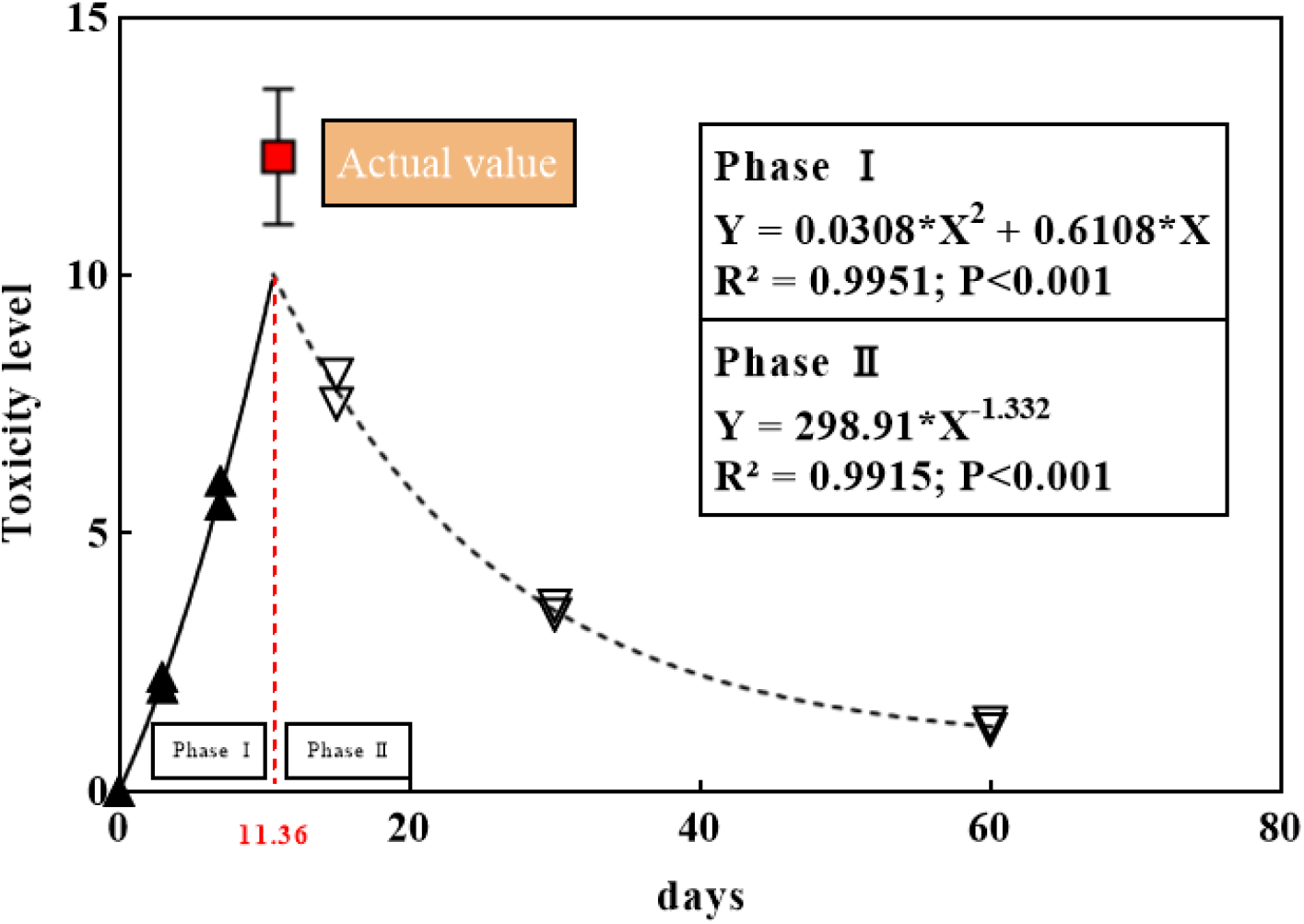
The dynamic of leachate toxicity over time. Two phases (I and II) were fitted with different functions. The intersection of two curves means the days (red number) corresponding to maximum leaching toxicity. The red box represents actual value in verification.

### 3.2 TWPs leachate toxicity cannot be characterized by single component

Among the 137 organic compounds identified in TWPs leachate using HPLC-MS, none exhibited a concentration-time trend that significantly correlated with overall leachate toxicity dynamic (**Figure 2**). Given the complex interactions among the hundreds of soluble substances in tires (Chen et al., 2024), it would be impractical to assess the biological toxicity of TWPs leachate solely based on specific components.

Based on the toxicity level-time functions shown in **Figure 2**, all nominal concentration series (250-1000 mg/L) at different extraction times were converted to doses within unified scale. Correlation of these doses with corresponding biological responses (BRI in **Figure 1**) enabled the generation of a standardized dose-response curve for the leachate (**Figure 3**). This curve conforms to the classical bell-shaped hormetic dose-response model (R^2^=0.7988; P<0.05), characterized by three distinct zones (I: lag zone; II: beneficial zone; III: adverse zone) and a maximum stimulatory response of 130%-160% relative to the control (Lushchak, 2014). This indicates that TWP leachate can be treated as a whole pollutant for analyzing its biological effects.

**Figure 3.**
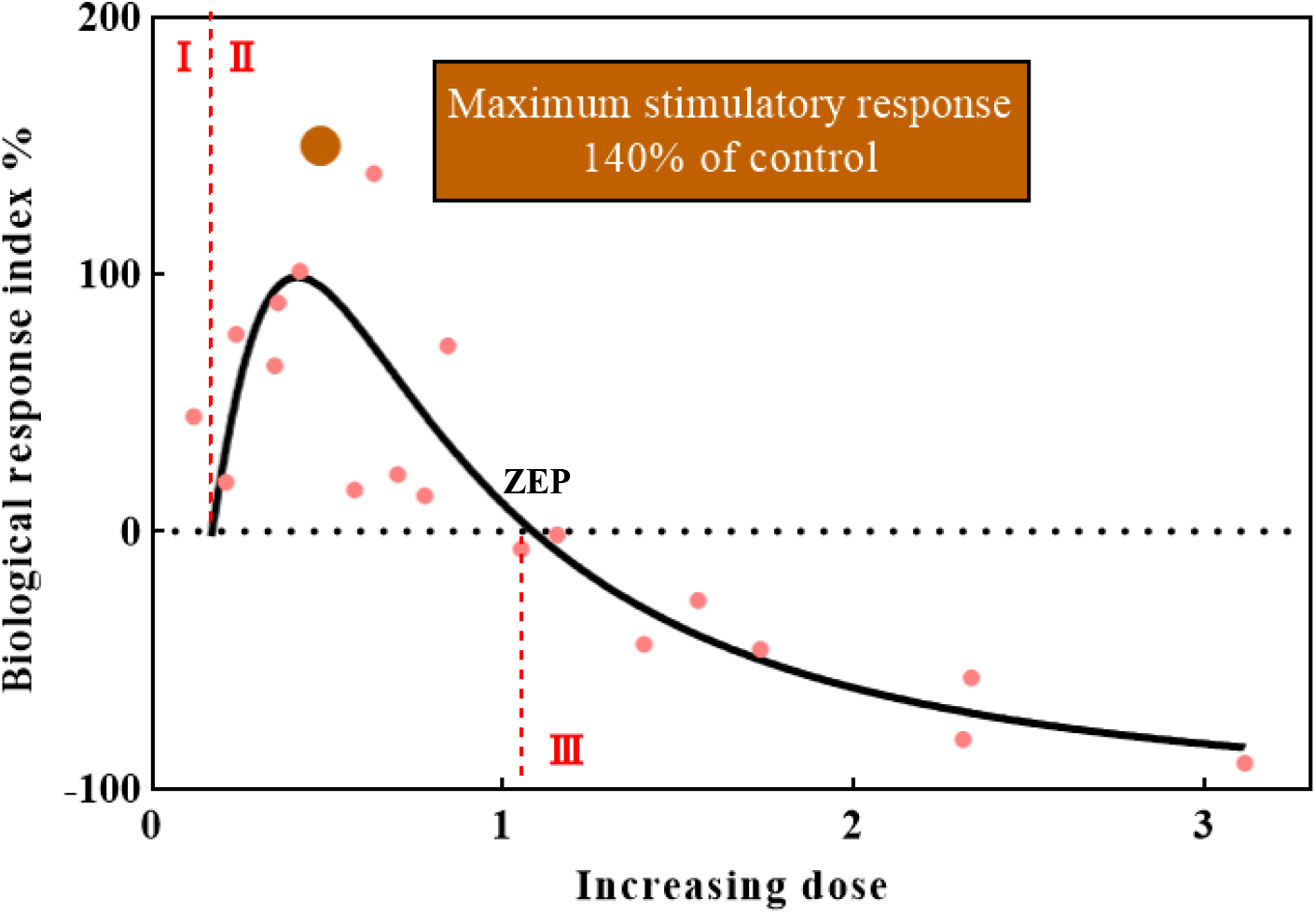
The dose-response curve of TWPs leachate. Biological response was reflected by BRI value. Positive value means hormesis, otherwise inhibition; I, II and III represent lag-zone, beneficial zone and adverse zone, and the bigger brown dot means the maximum hormetic effect value (140%). These characteristics are consistent with the typical hormesis dose-response model.

### 3.3 Leachate toxicity evolution of TWPs under in situ weathering conditions

Prior to the immersion of TWPs in water, UV irradiation and high temperature (70℃) had opposing effects on leachate toxicity, as shown in **Figure 4**. Compared to the control, high temperature increased leachate toxicity by up to 66%, while UV irradiation mitigated this increase by 84%. These results indicate an antagonistic interaction between UV irradiation and high temperature.

**Figure 4.**
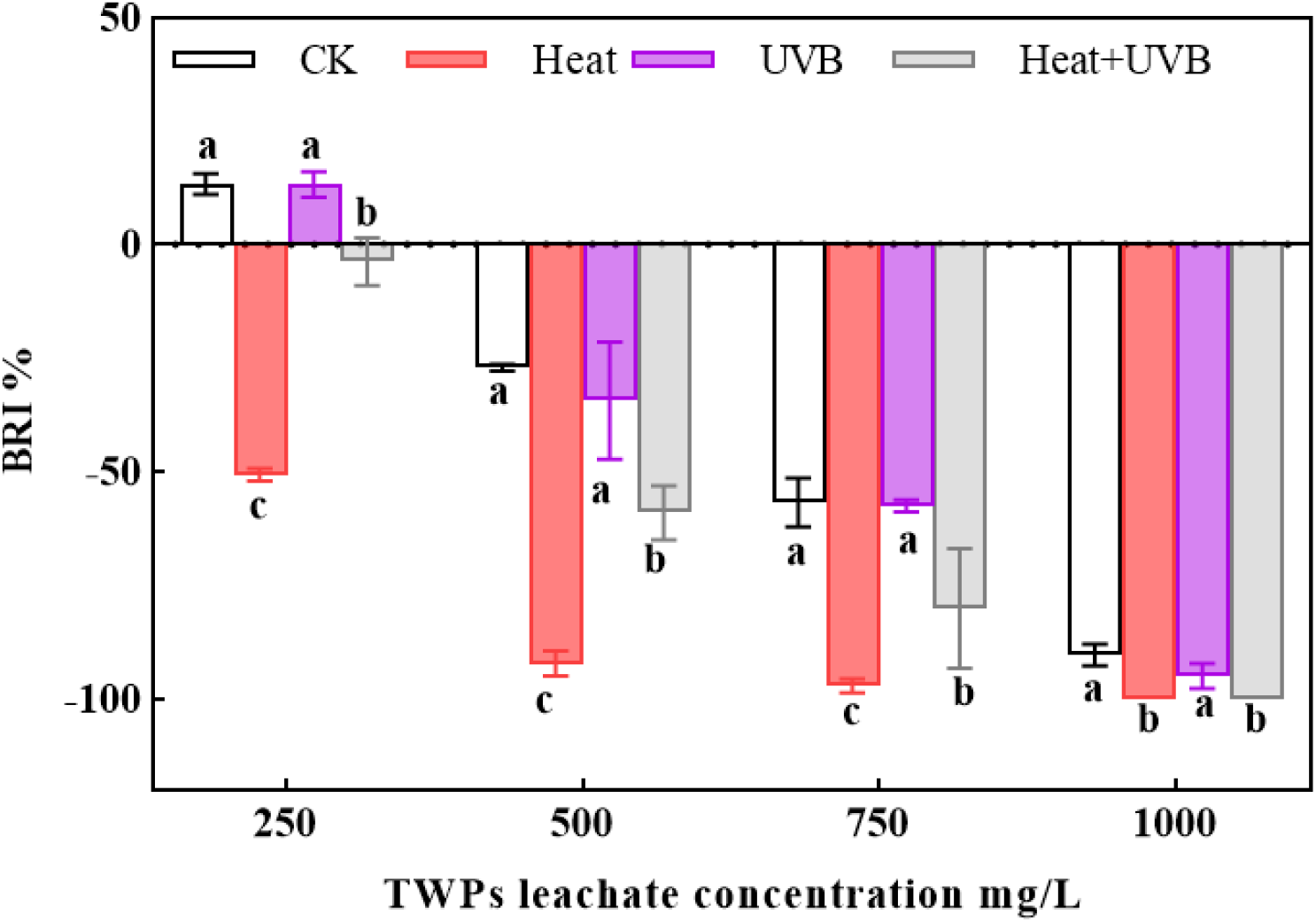
The biological responses to leaching toxicity from TWPs exposed to UV irradiation and high temperature. Positive BRI value means hermetic effect, otherwise inhibition. In each concentration group, the different lowercase letters mean significant differences (P<0.05).

The effects of high temperature and UV irradiation on leaching toxicity were attributed to changes in the concentration of leaching compounds (**Table 1**; Table S2). Compared to the control group, high temperature of 70℃ increased the concentration of 67 compounds by an average of 77%, including 26 compounds not detected in the control (**Table 1**, Heat-CK). Conversely, UV irradiation decreased the concentration of these compounds, effectively mitigating the effect of high temperature (**Table 1**, UV-Heat). These observed changes in leaching compounds align with the biological responses observed in the leachate. High temperature likely enhances the leaching process of toxic substances by increasing the mobility of bound toxicants in TWPs, as also noted in other studies (Kolomijeca et al., 2020; Lv et al., 2024). This is also supported by the enhanced volatilization observed at high temperatures (**Table 1**, Vol-CK). UV irradiation can induce photodegradation of specific toxic compounds in TWPs leachate via bond cleavage and functional group alteration (Fan et al., 2021; Kolomijeca et al., 2020; Zhang et al., 2024), likely mitigating the increased biological toxicity caused by high temperature.

**Table 1.** The changes of compounds concentration in TWPs leachate among different treatments.

| Treatment comparison | UP |  | DOWN |  |
| --- | --- | --- | --- | --- |
|  | Items (New) | Range | Items | Range |
| Heat-CK | 67 (26) | 77% | 48 | 42% |
| UV-CK | 53 (7) | 51% | 42 | 33% |
| UV-Heat | 21 | 40% | 97 | 69% |
| Vol-CK | 43 (6) | - | - | - |

**Notes:** CK, Heat and UV represent the treatments of control, 70°C high temperature for 12h and daily UVB irradiation dose. “Vol” means the liquid aired through gas during heating TWPs containing soluble volatile substances. The number in bracket means new compounds not detected in control group.

In this study, the exposure doses of UV irradiation and high temperature were comparable to those experienced in the environment over a single day. This suggests that the leachate toxicity is highly sensitive to these weathering conditions, even with daily removal of TWPs from urban roads. As UV irradiation and temperature exhibit temporal and spatial heterogeneity, the toxicity of TWP leachate will also show seasonal and regional fluctuations. Higher toxicity is expected in low latitude and altitude regions, as well as on cloudy summer days. Correspondingly, relevant policies, e.g., road cleaning frequency and timing, can be formulated to minimize the toxicity.

Moreover, compared to the control leachate, the volatile gas phase contained 43 water-soluble compounds at higher concentrations, out of a total of 87 identified compounds. (**Table 1**, Vol-CK). These compounds included benzene, its derivatives, and benzothiazoles, all of which are known for their high biological toxicity (Table S2). Additionally, several heavy metals were detected, including Cr (0.091 mg/L), Mn (0.068 mg/L), Co (0.009 mg/L), Ni (0.059 mg/L), Cu (0.024 mg/L), Zn (0.782 mg/L), and Cd (0.002 mg/L). Furthermore, the volume of volatile gas generated in this study was comparable to the average daily respiratory volume of an adult human. These components may be more bioavailable to humans, particularly residents of areas with high traffic density. This suggests that the volatile components, as a distinct and novel exposure pathway, warrant increased attention over the TWPs themselves. The chronic toxicity of these volatile components and their potential implications for human health demand further investigation.

## 4. Conclusions

The quantitative toxicological assessment of TWPs leachate is hindered by the ambiguity in dose represented by the amount of TWPs, as its leachate toxicity varies with leaching time and environmental factors. Correspondingly, this study presents the following key findings: 1) The kinetics of leachate toxicity over time is quantified based on biological responses, and characterized by biphasic model. 2) Assessing the toxicity of leachate through individual components is infeasible; the leachate should be treated as a complex mixture, as its dose-response curve aligns well with the classical hormesis model. 3) High temperature increases leachate toxicity by enhancing the release of toxic compounds, while UV irradiation mitigates this effect. 4) Volatile substances released under high temperature represent a distinct exposure pathway, independent of the TWPs themselves. These findings provide a framework for unifying and standardizing the assessment of TWPs leachate toxicity on a common scale.

## Acknowledgements

This work was supported by the Natural Science Foundation of Hainan Province (421QN199), Graduate Research and Innovation Project of Jiangsu Province (KYCX20_1190), Graduate Research and Innovation Project of Hainan (Qhys2021-216).

