## Supplemental Materials for "When and how leachate toxicity of tire wear particles peaks: quantifying its dynamics using dose-response analysis"

### Section 1: Material and methods

#### *Components analysis of leachate by LC-MS*

Chromatographic column ： C18 column (Zorbax Eclipse C18(1.8 μm*2.1mm*100mm)). Chromatographic separation conditions：The column temperature is 30°C; the flow rate is 0.3 mL/min；Mobile phase composition A：0.1% Fomic acid solution，B：pure acetonitrile；The injection volume： 2 μL，the autosampler temperature： 4 ℃.

Positive mode: heater temperature 325 °C; sheath gas flow: 45 arb（arbitrary units）; aux gas flow: 15 arb; sweep gas flow: 1 arb; electrospray voltage: 3.5 KV; capillary temperature: 330 °C; S-Lens RF Level: 55%.

Negative mode: heater temperature 325 °C; sheath gas flow: 45 arb; aux gas flow: 15 arb; sweep gas flow: 1 arb; electrospray voltage: 3.5 KV; capillary temperature: 330 °C; S-Lens RF Level: 55%.

Scanning mode: full scan (Full Scan, m/z 100~1500) and data-dependent mass spectrometry (dd-MS2, TopN = 10); resolution: 120,000 (MS1) & 60,000 (MS2). Collision Mode: High Energy Collision Dissociation (HCD).

**Table S1. LC mobile phase conditions**

| Time (min) | Flow rate (µL/min) | Gradient | B% Acetonitrile | A% Fomic acid |
|---|---|---|---|---|
| 0-2 | 300 | - | 5 | 95 |
| 2-6 | 300 | Linear gradient | 30 | 70 |
| 6-7 | 300 | - | 30 | 70 |
| 7-12 | 300 | Linear gradient | 78 | 22 |
| 12-14 | 300 | - | 78 | 22 |
| 14-17 | 300 | Linear gradient | 95 | 5 |
| 17-20 | 300 | - | 95 | 5 |
| 20-21 | 300 | Linear gradient | 5 | 95 |
| 21-25 | 300 | - | 5 | 95 |

### Section 2 Results

**Table S2. The change of components concentration in TWPs leachate among treatments**

| Name | Class | Heat-CK | UV-Heat | UV-CK | Vol-CK |
|---|---|---|---|---|---|
| Iminodimethanethiol | Thiohemiaminal derivatives | ↓ | — | ↓ | |
| Menadione bisulfite | Naphthalenes | ↓ | — | ↓ | |
| Aminomethanesulfonic acid | Organic sulfonic acids and derivatives | ↑ | ↓ | ↑ | ↑ |
| 4-Bromophenylboric acid | Benzene and substituted derivatives | ↑ | ↓ | ↑ | ↑ |
| 4-Nitrophenyl sulfate | Organic sulfuric acids and derivatives | ↑ | ↓ | ↑ | ↑ |
| 3-Bromo-4-hydroxybenzoic acid | Benzene and substituted derivatives | ↑ | ↓ | ↑ | ↑ |
| Tris(hydroxymethyl)aminomethane | Organonitrogen compounds | ↑ (New) | ↓ | — | |
| Diethanolamine | Organonitrogen compounds | ↓ | ↓ | ↓ | ↑ |
| Bicine | Carboxylic acids and derivatives | ↑ | ↓ | ↓ | ↑ |
| 2-(3-Iodo-2-propyn-1-yl)-2H-tetrazole | Azoles | ↓ | ↑ | ↑ | ↑ |
| Urocanic acid | Azoles | ↑ | ↓ | ↑ | ↑ |
| L-Pyroglutamic acid | Carboxylic acids and derivatives | — | — | — | |
| L-Leucine | Carboxylic acids and derivatives | — | — | — | |
| 4-Ethynylaniline | Benzene and substituted derivatives | ↑ (New) | ↓ | ↑ (New) | ↑ (New) |
| Caprolactam | Lactams | ↑ | ↓ | ↑ | ↑ |

| Name | Class | Heat-CK | UV-Heat | UV-CK | Vol-CK |
|---|---|---|---|---|---|
| PEG n6 | Organooxygen compounds | ↓ | ↑ | ↑ | ↑ |
| N,N-Diisopropylethylamine | Organonitrogen compounds | ↓ | ↓ | ↑ | ↑ |
| PEG n7 | Organooxygen compounds | — | ↑ | — | |
| Phthalic anhydride | Benzofurans | ↑ **(New)** | ↓ | — | |
| PEG n8 | Organooxygen compounds | — | ↑ | — | |
| Laurolactam | Macrolactams | ↑ | ↓ | ↓ | ↑ |
| N~2~-[4-({[3-(Cyclohexylamino)propyl]amino}methyl)benzyl]-6-(1-piperazinyl)-2,4-pyrimidinediamine | Diazinanes | ↓ | ↓ | ↓ | ↑ |
| Acetanilide | Benzene and substituted derivatives | — | — | — | ↑ **(New)** |
| Dicyclohexylamine | Organonitrogen compounds | ↑ | ↓ | ↑ | ↑ |
| N,N'-Bis[5-{[(3Z)-3-amino-3-(ethylimino)propyl]carbamoyl}-1-(3-methylbutyl)-1H-pyrrol-3-yl]terephthalamide | Benzene and substituted derivatives | ↑ | ↓ | ↑ | ↑ |
| Cyclo(D-leucyl-L-leucyl-L-leucyl-L-leucyl-L-leucyl-L-leucyl) | Carboxylic acids and derivatives | ↑ | ↓ | ↑ | ↑ |
| Diphenylphosphinic acid | Benzene and substituted derivatives | — | — | — | |
| (2S,3R)-2-Amino-1,3-dodecanediol | Organonitrogen compounds | ↓ | ↓ | ↓ | ↑ |

| Name | Class | Heat-CK | UV-Heat | UV-CK | Vol-CK |
|---|---|---|---|---|---|
| 2-Hydroxybenzothiazole | Benzothiazoles | ↑ **(New)** | ↓ | — | ↑ **(New)** |
| 4,6-Di-t-butylpyrogallol | Phenols | ↑ **(New)** | ↓ | — | |
| N-(2,4-Dimethylphenyl)formamide | Benzene and substituted derivatives | ↑ | ↓ | ↓ | ↑ |
| Benzothiazole | Benzothiazoles | — | — | — | |
| 3-Hydroxytetradecanedioic acid | Fatty Acyls | ↑ **(New)** | ↓ | — | |
| 2,6-Di-tert-butyl-1,4-benzoquinone | Prenol lipids | ↑ **(New)** | ↓ | — | |
| Betaxolol | Phenols | ↓ | — | ↑ | |
| 3-(Benzylamino)-5-(4-chlorophenyl)cyclohex-2-en-1-one | Benzene and substituted derivatives | — | — | — | |
| 1-[(2-Hydroxyethyl)amino]-2-dodecanol | Organonitrogen compounds | ↓ | ↓ | ↓ | ↑ |
| N,N-Dimethyldecylamine N-oxide | Organonitrogen compounds | ↓ | ↓ | ↓ | ↑ |
| Harmine | Harmala alkaloids | ↓ | ↓ | ↓ | ↑ |
| Metyrapone | Organooxygen compounds | ↓ | ↑ | ↑ | ↑ |
| N-dodecyl ethanolamine | Organonitrogen compounds | ↑ | ↓ | ↓ | ↑ |
| 13(S)-HOTrE | Fatty Acyls | ↑ **(New)** | ↓ | — | |
| Phthalic acid | Benzene and substituted derivatives | ↑ **(New)** | ↓ | — | |
| 2-Amino-1,3,4-octadecanetriol | Organonitrogen compounds | ↓ | ↓ | ↓ | ↑ |
| Monobutyl phthalate | Benzene and substituted derivatives | ↑ **(New)** | ↓ | — | |

| **Name** | **Class** | **Heat-CK** | **UV-Heat** | **UV-CK** | **Vol-CK** |
|---|---|---|---|---|---|
| Lauryldimethylamine oxide | Organonitrogen compounds | ↓ | ↓ | ↓ | ↑ |
| Hydrolyzed fumonisin B1 | Organonitrogen compounds | ↓ | ↓ | ↓ | |
| 2-Amino-1,3,4,5-icosanetetrol | Organonitrogen compounds | ↓ | ↓ | ↓ | ↑ |
| C16 phytosphingosine | Organonitrogen compounds | ↑ | ↓ | ↑ | ↑ |
| Dodecylamine | Organonitrogen compounds | ↓ | ↓ | ↓ | ↑ |
| N,N'-Dicyclohexylurea | Organic carbonic acids and derivatives | ↓ | ↓ | ↓ | |
| 1-Tetradecylamine | Organonitrogen compounds | ↓ | ↓ | ↑ | ↑ |
| 11β-Hydroxyandrosterone | Steroids and steroid derivatives | ↑ **（New）** | ↓ | — | |
| AUDA | Fatty Acyls | ↑ | ↓ | ↑ | ↑ |
| Bis(methylbenzylidene)sorbitol | Cinnamyl alcohols | ↑ | ↓ | ↓ | ↑ |
| Safingol | Organonitrogen compounds | ↓ | ↓ | ↓ | ↑ |
| C20 phytosphingosine | Organonitrogen compounds | ↓ | ↓ | ↓ | ↑ |
| MDPBP | Benzene and substituted derivatives | ↓ | — | ↓ | ↑ |
| Myristamine oxide | Organonitrogen compounds | ↓ | ↓ | ↓ | ↑ |
| 2-Methyl-S-benzothiazole | Benzothiazoles | ↓ | ↓ | ↓ | ↑ |
| Bis(4-ethylbenzylidene)sorbitol | Dioxanes | ↑ | ↓ | ↑ | ↑ |
| N,N-Bis(2-hydroxyethyl)dodecanamide | Fatty Acyls | ↑ | ↓ | ↓ | ↑ |

| Name | Class | Heat-CK | UV-Heat | UV-CK | Vol-CK |
|---|---|---|---|---|---|
| Diphenylamine | Benzene and substituted derivatives | ↓ | — | ↓ | |
| 4-Methylbenzophenone | Benzene and substituted derivatives | — | ↑ | — | |
| Cetrimonium | Organonitrogen compounds | ↑ (New) | ↓ | ↑ (New) | ↑ (New) |
| 2,3-Methylenedioxy pyrovalerone | Benzene and substituted derivatives | ↑ | ↓ | ↑ | ↑ |
| 4-Chlorobenzophenone | Benzene and substituted derivatives | ↑ (New) | ↓ | — | |
| 2-hydroxyimipramine | Benzazepines | ↓ | ↑ | ↑ | ↑ |
| Tradecamide | Fatty Acyls | ↑ | ↓ | ↑ | ↑ |
| butyrin | Glycerolipids | ↑ | ↓ | ↑ | ↑ |
| 1,5,9-Trimethyl-11,14,15,16-tetraoxatetracyclo[10.3.1.0~4,13~.0~8,13~]hexadecan-10-ol | Prenol lipids | ↑ | ↓ | ↑ | ↑ |
| N-Cyclohexyl-1-(10,11-dihydro-5H-dibenzo[a,d][7]annulen-5-yl)-3-azetidinamine | Dibenzocycloheptenes | ↓ | ↑ | ↓ | |
| Leucomalachite green | Triphenyl compounds | — | — | — | |
| α-Linolenoyl ethanolamide | Organonitrogen compounds | ↓ | ↑ | ↑ | ↑ |
| Dodecamethylcyclohexasiloxane | Organometalloid compounds | — | — | — | |
| 3-Hydroxy-2-methylpyridine | Pyridines and derivatives | ↑ (New) | ↓ | — | |
| Methyl isonicotinate | Pyridines and derivatives | ↑ (New) | ↓ | — | |

| Name | Class | Heat-CK | UV-Heat | UV-CK | Vol-CK |
|---|---|---|---|---|---|
| N,N-Dipentyl-4-(2-(4-quinolinyl)vinyl)aniline | Quinolines and derivatives | — | — | — | |
| Oleamide | Fatty Acyls | — | ↑ | — | |
| Myristoyl glutamic acid | Carboxylic acids and derivatives | — | ↑ | — | |
| 2-{4-[1-(4-Fluoro-2-methylphenyl)-4-piperidinyl]-1-methyl-2-piperazinyl}ethanol | Piperidines | — | ↑ | — | |
| 1-(2-Ethoxyphenyl)-3-[2-(4-propyl-1-piperazinyl)ethyl]urea | Benzene and substituted derivatives | — | ↑ | — | |
| Stearamide | Carboximidic acids and derivatives | ↓ | ↓ | ↑ | ↑ |
| Molybdenite | Transition metal organides | ↑ | ↓ | ↑ | ↑ |
| 3,4-Dichloro-5-[(3-cyano-4-fluorophenyl)sulfamoyl]benzoic acid | Benzene and substituted derivatives | ↑ | ↓ | ↑ | ↑ |
| Ethyl N-[(5-bromo-2-thienyl)carbonyl]glycinate | Carboxylic acids and derivatives | ↑ | ↓ | ↑ | ↑ |
| 2,6-Dichlorotoluene | Benzene and substituted derivatives | ↑ | ↓ | ↑ | ↑ |
| 4-[(Trifluoromethyl)sulfanyl]phenyl dihydrogen phosphate | Organic phosphoric acids and derivatives | ↑ | ↓ | ↑ | ↑ |
| 2-[(5-Bromo-1,3-thiazol-2-yl)amino]-2-oxoethyl (4-chlorophenoxy)acetate | Peptidomimetics | ↑ | ↓ | ↑ | ↑ |
| 2,5-Dichloro-4-nitro-N-[4-(trifluoromethyl)phenyl]-3-thiophenesulfonamide | Benzene and substituted derivatives | ↑ | ↓ | ↓ | ↑ |

| Name | Class | Heat-CK | UV-Heat | UV-CK | Vol-CK |
|---|---|---|---|---|---|
| 2,6-Difluoro-N-(3-thioxo-3H-1,2,4-dithiazol-5-yl)benzamide | Benzene and substituted derivatives | ↑ | ↓ | ↑ | ↑ |
| (2Z)-3-(3-Iodophenyl)-2-sulfanylacrylic acid | Cinnamic acids and derivatives | ↓ | ↓ | ↓ | ↑ |
| (4-Bromophenyl)[2-(methylamino)-5-thiazolyl]methanone | Organooxygen compounds | ↑ | ↓ | ↑ | ↑ |
| 5-Iodo-1H-indazole | Benzopyrazoles | ↓ | ↓ | ↓ | ↑ |
| 2,4,6-Trichlorophenyl 2-phenylethanesulfonate | Benzene and substituted derivatives | ↑ | ↓ | ↑ | ↑ |
| (3,4-Dihydroxy-5-iodobenzylidene)malononitrile | Phenols | ↑ | ↓ | ↑ | ↑ |
| 12,13-Dibromo-2,3,5,6,8,9-hexahydro-1,4,7,10-benzotetraoxacyclododecine | Organooxygen compounds | ↓ | ↑ | ↓ | ↑ |
| (3Z,5Z)-3,5-Bis(3-bromobenzylidene)tetrahydro-4H-thiopyran-4-one | Benzene and substituted derivatives | ↑ | ↓ | ↓ | ↑ |
| N~2~-[(2,5-Dibromophenyl)sulfonyl]-N-(1,1-dioxidotetrahydro-3-thiophenyl)-N~2~-ethylglycinamide | Benzene and substituted derivatives | ↓ | ↑ | ↓ | ↑ |
| 5-Chloro-4-nitro-N-[4-(trifluoromethyl)phenyl]-2-thiophenesulfonamide | Benzene and substituted derivatives | ↓ | ↑ | ↑ | ↑ |
| Mitotane | Benzene and substituted derivatives | — | — | ↑ **(New)** | |
| 2,3-Bis(4-bromophenyl)-4(3H)-quinazolinone | Diazanaphthalenes | ↓ | ↑ | ↑ | ↑ |
| Ethyl 3-(4-hydroxy-3,5-diiodophenyl)-2-phenylpropanoate | Stilbenes | ↑ **(New)** | ↓ | ↑ **(New)** | |

| Name | Class | Heat-CK | UV-Heat | UV-CK | Vol-CK |
|---|---|---|---|---|---|
| 7-methylbenzopentathiepin | Others | ↑ | ↓ | ↓ | ↑ |
| (2E)-3-[3-(Benzyloxy)phenyl]-1-(3-bromophenyl)-2-propen-1-one | Linear 1,3-diarylpropanoids | ↑ | ↓ | ↑ | ↑ |
| Citric acid | Carboxylic acids and derivatives | ↓ | ↓ | ↓ | ↑ |
| dimethyl citrate | Carboxylic acids and derivatives | ↑ | ↓ | ↑ | ↑ |
| N-Acetyl-D-alloisoleucine | Carboxylic acids and derivatives | ↑ **（New）** | ↓ | ↑ **（New）** | ↑ **（New）** |
| Isophthalic acid | Benzene and substituted derivatives | ↑ **（New）** | ↓ | ↑ **（New）** | |
| 8-Amino-7-oxononanoic acid | Fatty Acyls | ↓ | ↓ | ↓ | ↑ |
| N-acetyl-L-leucyl-N-{(2R)-6-[(tert-butoxycarbonyl)amino]-1-oxohexan-2-yl}-D-leucinamide | Carboxylic acids and derivatives | ↓ | ↓ | ↓ | ↑ |
| N-{[(1R)-2-{[(6R)-8-hydroxy-6,11-dimethyl-1,4,5,6-tetrahydro-2,6-methano-3-benzazocin-3(2H)-yl]methyl}-1-phenylcyclopropyl]carbonyl}glycyl-N-[6-(propanimidoylamino)hexyl]leucinamide | Carboxylic acids and derivatives | ↓ | ↑ | ↑ | ↑ |
| Cyclo(D-leucyl-L-leucyl-L-leucyl-L-leucyl-L-leucyl-L-leucyl) | Carboxylic acids and derivatives | ↑ **（New）** | ↓ | — | |
| Methyl N-methyl-N-(7-octenoyl)-L-valyl-N-methyl-L-valyl-N-methyl-L-valyl-N-methyl-L-alanyl-N-methyl-L-phenylalaninate | Carboxylic acids and derivatives | ↑ | ↓ | ↑ | ↑ |

| Name | Class | Heat-CK | UV-Heat | UV-CK | Vol-CK |
|---|---|---|---|---|---|
| (2R)-1-(beta-D-Galactopyranosyloxy)-3-(tetradecanoyloxy)-2-propanyl (9E,12E,15E)-9,12,15-octadecatrienoate | Others | ↓ | ↑ | ↑ | ↑ |
| Clorophene | Benzene and substituted derivatives | — | — | — | |
| Butyldiglycol acetate | Carboxylic acids and derivatives | ↑ **(New)** | ↓ | — | |
| Trinexapac | Vinylogous acids | — | — | — | |
| 1,2-Benzisothiazolin-3-one | Benzothiazoles | ↑ **(New)** | ↓ | — | ↑ **(New)** |
| 3-Hydroxytetradecanedioic acid | Fatty Acyls | ↑ **(New)** | ↓ | — | |
| MFCD00054545 | Organooxygen compounds | ↑ **(New)** | ↓ | — | |
| N-(1-{1-[(1-Ethyl-1H-pyrazol-4-yl)methyl]-4-piperidinyl}-1H-pyrazol-5-yl)cyclopentanecarboxamide | Organonitrogen compounds | ↓ | ↓ | ↑ | ↑ |
| Dodecanedioic acid | Fatty Acyls | ↑ **(New)** | ↓ | — | |
| Tetradecanedioic acid | Fatty Acyls | ↑ **(New)** | ↓ | — | |
| Monobutyl phthalate | Benzene and substituted derivatives | ↑ | ↓ | ↑ | ↑ |
| (15Z)-9,12,13-Trihydroxy-15-octadecenoic acid | Fatty Acyls | ↑ **(New)** | ↓ | — | |
| N,N-Bis(2-hydroxyethyl)dodecanamide | Fatty Acyls | ↓ | ↓ | ↓ | ↑ |
| 2,5-di-tert-Butylhydroquinone | Benzene and substituted derivatives | ↓ | ↓ | ↓ | ↑ |
| Butylparaben | Benzene and substituted derivatives | ↓ | — | ↓ | |

| Name | Class | Heat-CK | UV-Heat | UV-CK | Vol-CK |
|---|---|---|---|---|---|
| Amipylo | Carboxylic acids and derivatives | ↑ | ↓ | ↑ | ↑ |
| 2,2'-Methylenebis(4-methyl-6-tert-butylphenol) | Benzene and substituted derivatives | ↓ | ↓ | ↑ | ↑ |
| 1-(2-Ethoxyphenyl)-3-[2-(4-propyl-1-piperazinyl)ethyl]urea | Benzene and substituted derivatives | — | ↑ | — | |
| (3R,4S)-3-{(1R)-1-Acetamido-2-[(1-methoxy-2-butanyl)amino]-2-oxoethyl}-4-[(diaminomethylene)amino]cyclopentanecarboxylic acid | Carboxylic acids and derivatives | — | ↑ | — | |
| palmitic acid | Fatty Acyls | ↓ | ↓ | ↓ | ↑ |

**Notes: CK, Heat and UV represent the treatments of control, 70℃ high temperature for 12h and daily UVB irradiation dose. Vol means the liquid aired through gas during heating TWPs containing soluble volatile substances. ↑: Increasing, ↓: Reducing, —: No change, (New): New compounds not detected in control.**
